# Microextrusion Printing Cell-Laden Networks of Type I Collagen with Patterned Anisotropy and Geometry

**DOI:** 10.1101/509265

**Authors:** Bryan A. Nerger, P.-T. Brun, Celeste M. Nelson

## Abstract

Type I collagen self-assembles into three-dimensional (3D) fibrous networks. These dynamic viscoelastic materials can be remodeled in response to mechanical and chemical cues to form anisotropic networks, the structure of which influences tissue development, homeostasis, and disease progression. Conventional approaches for fabricating anisotropic networks of type I collagen are often limited to unidirectional alignment over small areas. Here, we describe a new approach for engineering cell-laden anisotropic networks of type I collagen fibers using 3D microextrusion printing of a collagen-Matrigel ink. By adding molecular crowders, we demonstrate hierarchical control of 3D-printed collagen with the ability to spatially pattern collagen fiber anisotropy and geometry. Our data suggest that collagen anisotropy results from a combination of molecular crowding in the ink and shear and extensional flows present during 3D-printing. We demonstrate that human breast cancer cells cultured on 3D-printed collagen orient along the direction of collagen fiber alignment. We also demonstrate the ability to simultaneously bioprint epithelial cell clusters and control the alignment and geometry of collagen fibers surrounding cells in the bioink. The resulting cell-laden constructs consist of epithelial cell clusters fully embedded in aligned networks of collagen fibers. We foresee that cell-laden collagen-Matrigel constructs with spatially-patterned anisotropy and geometry will be broadly useful for the fields of developmental biology, tissue engineering, and regenerative medicine.

## Introduction

The extracellular matrix (ECM) consists of a heterogeneous mixture of macromolecules that form the non-cellular component of tissues ^1^. The fibrous structure of the ECM serves as a scaffold that provides mechanical support and chemical cues to its constituent cells. Reciprocal biochemical and biophysical interactions lead to continuous remodeling of the ECM, giving rise to a dynamic viscoelastic material with a rich diversity of structures and functions. One of the most common ECM structural motifs consists of anisotropic networks of type I collagen, which are associated with biological processes as diverse as collective cell migration ^2^, wound healing ^3^, metastasis ^4^, and tissue morphogenesis ^5^. These anisotropic patterns of collagen fibers have been challenging to recapitulate *ex vivo*.

Collagen fibers are formed *in vitro* by the self-assembly of 300-nm-long tropocollagen monomers ^6^. Self-assembly is driven by a large positive entropy that results from the displacement of structured water surrounding tropocollagen monomers ^7, 8^, and depends on several parameters including concentration, pH, temperature, ionic strength, and molecular crowding ^9-11^. The resulting networks recapitulate key mechanical and chemical features observed *in vivo* and have been used extensively as models for native ECM ^1^.

Nonetheless, *in vitro* networks of collagen are heterogeneous and lack the anisotropy observed *in vivo*. In response to these limitations, several approaches have been described to alter collagen anisotropy either during or after self-assembly *in vitro*. During self-assembly, collagen fiber anisotropy can be influenced by magnetic fields ^12^, flow fields ^13-15^, shear ^16, 17^, mechanical instabilities ^18^, molding ^19^, or a combination of molecular crowding and spatial confinement ^20^, among others. After self-assembly, collagen fiber anisotropy can be generated by applying mechanical strain ^21, 22^ or shear ^16^. To the best of our knowledge, these approaches can only align collagen fibers uniaxially and are not able to spatially pattern collagen fiber orientation and geometry. As a result, existing approaches are unable to reproduce the complexity of collagen structures in native ECM. Moreover, it is challenging to incorporate fully embedded cells or tissues into aligned networks of collagen using existing techniques. While strained PDMS molds can incorporate cells into networks of collagen with aligned fibers, it is unclear how to isolate the effects of fiber alignment, compression, and collagen densification on cell behavior ^23^.

Here, we describe a new approach to fabricate cell-laden anisotropic networks of type I collagen fibers using 3D microextrusion printing of collagen-Matrigel inks. We show that collagen can be 3D-printed while simultaneously controlling the spatial deposition, geometry, and anisotropy of the resulting fibrous network. While numerous studies have 3D-printed collagen inks ^24-28^, we incorporate Matrigel into a collagen ink in order to print significantly lower collagen concentrations (0.8 mg/ml). Our approach allows the collagen fiber geometry and orientation to be precisely analyzed throughout the volume of the printed construct. In addition, the low concentration of collagen more accurately reproduces the networks of collagen fibers used in cell culture experiments in studies of cancer cell migration ^29^ and tissue-ECM interactions ^30^. We demonstrate that molecular crowding in a collagen ink as well as substratum hydrophobicity can be used to tune collagen fiber alignment. We also demonstrate the ability to generate collagen fiber networks with multidirectional collagen fiber alignment. By combining 3D microextrusion printing and drop-casting, we show that the collagen fiber geometry and alignment can be patterned over mm-length scales, wherein distinct collagen fiber morphologies are separated by sharp interfaces with tunable geometry. Lastly, we show for the first time that microextrusion printing can be used to simultaneously bioprint epithelial cell clusters and control the alignment and geometry of collagen fibers surrounding these fully-embedded clusters. Compared to other approaches, 3D microextrusion printing of collagen-Matrigel hydrogels is a simple, fast, and versatile technique that can generate large-scale cell-laden constructs with spatial control of collagen fiber alignment and geometry.

## Materials and Methods

### Reagents and collagen gel preparation

Acid-solubilized bovine type I collagen (Advanced Biomatrix, Carlsbad, CA) was adjusted to a final concentration of 0.8 mg/ml and neutralized to pH∼8 with the manufacturer-provided neutralizing solution. Laponite XLG (BYK Additive and Instruments, Gonzales, TX), Pluronic F127 (BASF, Florham Park, NJ), or Matrigel (Corning, Corning, NY) were used as gelatinous additives at concentrations of 3 mg/ml, 250 mg/ml, and 4.2-10.1 mg/ml, respectively. The protein concentration of Matrigel was adjusted by dilution with 1:1 DMEM:F12 medium (Life Technologies, Carlsbad, CA). The molecular crowders, Ficoll 70 and Ficoll 400 (Sigma-Aldrich), were first dissolved in phosphate-buffered saline and used at final concentrations of 6, 12, 18, and 24 mg/ml. The pH-adjusted collagen mixture was pipetted into a 3D-printing syringe and stored at ∼0°C for 1 h before printing.

### 3D microextrusion printing

Collagen inks were 3D-printed using a microextrusion bioprinter (Inkredible+, CELLINK, Sweden) and conical polyethylene nozzles with diameters of 200 µm, 254 µm, and 400 µm (Nordson EFD, Robbinsville, NJ). All inks were 3D-printed at room temperature (∼20°C) and were stored for ∼5 min at room temperature prior to printing. The printing pressure and speed varied from 1-40 kPa and 20-80 mm/s, respectively. Printing paths were generated by writing G-code or by drawing 3D objects in Autodesk Inventor (Autodesk, San Rafael, CA) and importing the resulting STL file into Slic3r to generate G-code. Samples were 3D-printed onto a no. 1 glass coverslip at room temperature and cured in a 37°C incubator at 5% CO_2_ for 1 h before imaging. Prior to printing, coverslips were rinsed with 100% ethanol, air dried, and then cleaned with a UV/ozone cleaner (Jelight Company, Irvine, CA) for 7 min. Clean coverslips were silanized by exposure to trichloro(1H,1H,2H,2H-perfluorooctyl)silane (TCPFOS) (Sigma-Aldrich) vapors under vacuum for 24 hours, 3,3,3-trifluoropropyl-trichlorosilane (TFPTCS) (Alfa Aesar, Haverhill, MA) vapors under vacuum for 20 min, or trimethylchlorosilane (TMCS) (Sigma-Aldrich) vapors at atmospheric pressure for 20 min. Treatment with methoxy-poly(ethylene glycol)-silane (PEG-silane) (Laysan Bio Inc, Arab, AL) was achieved by coating clean coverslips with an ethanolic solution of 0.5% PEG-silane and 1% acetic acid for 30 min at 70°C.

### Cell culture

MDA-MB-231 human breast cancer cells were cultured using 1:1 DMEM:F12 medium (Life Technologies) supplemented with 50 µg/ml gentamicin (Sigma-Aldrich) and 10% fetal bovine serum (Atlanta Biologicals, Flowery Branch, GA). Normal EpH4 mammary epithelial cells were cultured using 1:1 DMEM:F12 medium (Life Technologies) supplemented with 50 µg/ml gentamicin (Sigma-Aldrich), 2% fetal bovine serum (Atlanta Biologicals, Flowery Branch, GA), and 5 µg/ml insulin (Sigma-Aldrich). Both cell lines were cultured in an incubator maintained at 37°C and 5% CO_2_. Clusters of mammary epithelial cells were generated by suspending cells in culture medium supplemented with 0.1% (w/v) Pluronic F108 (BASF, Ludwigshafen, Germany) and incubating at 37°C and 5% CO_2_ overnight. The culture medium for mammary epithelial cell clusters was supplemented with 5 ng/ml hepatocyte growth factor (HGF) (Sigma-Aldrich).

### Microscopy

Collagen fibers were imaged using a 10× or 20× air objective or a 40× oil-immersion objective and a Nikon A1 laser-scanning confocal microscope in reflection mode (488 nm argon laser with GaAsP detector). 30-µm z-stacks, scanned at 2-µm intervals, were acquired for each sample and the maximum-intensity z-projection was obtained using ImageJ (NIH). Confocal reflection microscopy (CRM) was also used to image collagen fiber orientation at 10-µm intervals throughout the depth of the sample. Cells cultured with 3D-printed collagen inks were imaged using a Nikon Plan Fluor 10×/0.30 *NA* or 20×/0.45 *NA* air objective and ORCA-03G digital CCD camera (Hamamatsu Photonics, Japan).

### Quantifying collagen fiber alignment and geometry

Collagen fiber alignment was quantified using the local gradient orientation method (created by Jean-Yves Tinevez) in ImageJ. Collagen fiber anisotropy was estimated using an alignment fraction, which represents the fraction of aligned intensity gradients with respect to the total number of identified intensity gradients in an image. We considered an intensity gradient oriented within 20° of the printing direction to be aligned. All fiber alignment calculations were performed using maximum-intensity z-projections of 30-µm z-stacks obtained using CRM. Lengths of collagen fiber bundles were measured manually using ImageJ, while fiber diameters were approximated using the BoneJ plug-in (created by Michael Doube) in ImageJ ^31^. Heat maps were generated using the heatmap function in MATLAB (R2015b; MathWorks, Natick, MA).

### Rheological measurements

Experiments were conducted using an MCR501 stress-controlled rheometer (Anton Paar, Ashland, VA). The temperature-dependent shear loss (G′′) and storage (G′) moduli were measured using a 25-mm parallel-plate with a 600-µm gap. Before testing, samples (∼300 µl) were loaded onto the bottom plate at a temperature of 4°C. The temperature was raised to 37°C at the beginning of the measurement to initiate gelation, and samples were oscillated at 1 rad/s and 0.5% strain for 10 min. Three independent measurements were acquired, and the average steady-state moduli were used for comparison.

### Contact angle measurements

Advancing and receding contact angles were measured at room temperature using a Model 500-F1 contact angle goniometer (ramé-hart, Succasunna, NJ). All measurements were conducted on glass substratum and reported values represent an average of 10 measurements. Collagen and collagen-Matrigel samples were neutralized immediately before contact angle measurements.

### Statistical analysis

Unless stated otherwise, data represent the mean of three independent replicates and error bars represent the standard error of the mean. Each independent replicate was conducted in triplicate, and two measurements were acquired for each sample. Statistical comparisons were performed using one-sided or two-sided *p*-values, which were calculated using Welch’s *t*-test or one-way ANOVA. A *p*-value less than 0.05 was considered to be statistically significant.

## Results and discussion

We began by evaluating the printability and shape retention of solutions of type I collagen during 3D microextrusion printing. “Shape retention” is a binary metric that describes the ability of a material to retain its shape after 3D-printing. Materials with acceptable shape retention should demonstrate less than 10% change in length and width upon reaching equilibrium as compared to the programmed dimensions of the object. “Printability” is a qualitative description of the ability of a material to be continuously extruded at a constant printing speed and pressure and serves as a binary metric to assess the feasibility of an ink for microextrusion printing. Inadequate printability can be caused by clogging of or inconsistent flow through the nozzle.

A neutralized collagen solution was pre-incubated on ice for 1 h prior to drop casting or 3D-printing (**Figure 1a**). Low-temperature pre-incubation allows collagen self-assembly to initiate at a reduced rate^32^. Confocal reflection microscopy (CRM) revealed that both drop-cast (**Figure 1b**) and 3D-printed (**Figure 1c**) collagen inks form isotropic collagen fiber networks. Moreover, we found that the collagen ink had inadequate shape retention (**Figure S1**) and could not be accurately 3D-printed into rectangular geometries. To enhance shape retention without increasing collagen concentration, rheological improvements are required.

**Figure 1.**
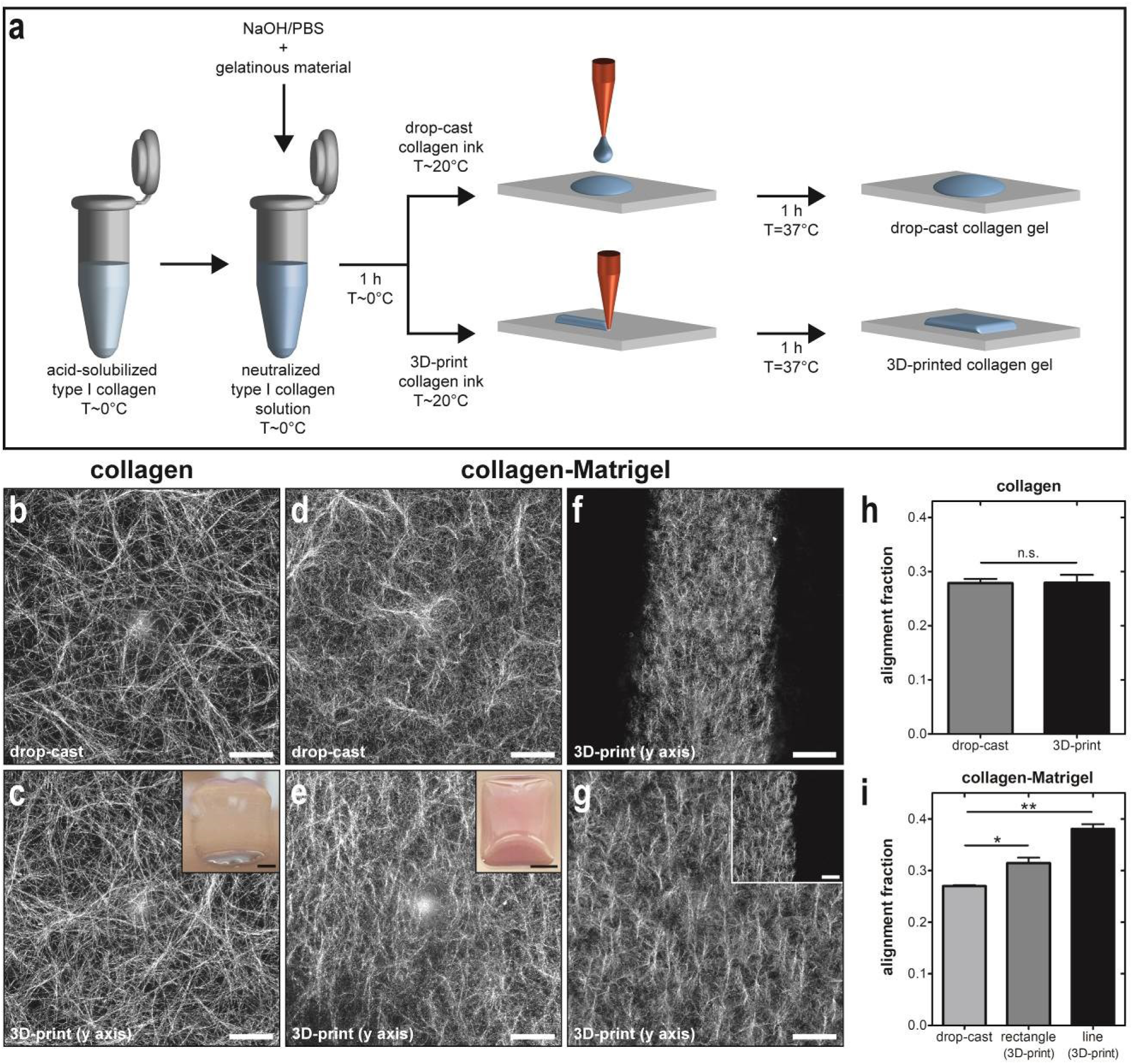
Designing an ink for 3D microextrusion printing of type I collagen. a) Schematic depicting ink formulation, drop-casting, and 3D-printing. CRM images of b) drop-cast and c) 3D-printed collagen and d) drop-cast and e) 3D-printed collagen-Matrigel. Scale bars on CRM and optical images represent 50 µm and 2.5 mm, respectively. Insets show representative 3D-printed rectangles. f) CRM image of 3D-printed line of collagen-Matrigel. Scale bar = 200 µm. g) Higher magnification CRM image of 3D-printed line (inset shows edge of printed line at the same magnification; scale bars = 100 µm). Average alignment fraction in drop-cast and 3D-printed samples of h) collagen and i) collagen-Matrigel. An average of 7 printed lines was used for quantification. All samples were 3D-printed onto no. 1 glass coverslips using 254-µm-diameter conical nozzles at a printing speed of 40 mm/s. All CRM images represent the maximum-intensity z-projection of a 30-µm z-stack. A Matrigel concentration of ∼5.0 mg/ml was used for collagen-Matrigel samples.

Shear-thinning materials, including physically crosslinked gels, are commonly used to improve the rheological properties of 3D-printing inks ^33, 34^. Shear and extensional flows generated during extrusion temporarily disrupt physical crosslinks, allowing a material to flow. After exiting the nozzle, the physical crosslinks reform and allow the printed shape to be retained. We therefore proceeded to mix collagen with shear-thinning gels to determine whether this would improve shape retention.

Aqueous Laponite and Pluronic F127, which have well-characterized shear-thinning properties, have been used extensively as rheological modifiers for 3D-printing and commercial applications ^35, 36^. Laponite is a synthetic hectorite clay that consists of nm-scale platelets. Pluronic F127 is a triblock copolymer of polyethylene oxide (PEO) and polypropylene oxide (PPO) (PEO-PPO-PEO), which forms micelles when dissolved in water above its critical micelle concentration (21 w/w%) under ambient conditions ^37^. Previous studies have suggested that Laponite ^38^ and Pluronic ^39, 40^ might be compatible additives to improve the mechanical properties of collagen.

We combined Laponite or Pluronic F127 with neutralized type I collagen and investigated their effects on shape retention, printability, and network structure. We found that collagen-Laponite inks, which consisted of collagen mixed with a 3 mg/ml Laponite gel, frequently clogged the nozzle during printing (**Figure S1**). In parallel, we mixed collagen with a 250 mg/ml solution of Pluronic F127 and observed that collagen-Pluronic inks frequently clogged the nozzle and did not retain their shape after printing (**Figure S1**).

To understand why Laponite and Pluronic F127 adversely impact collagen shape retention and printability, we imaged gels using CRM, which revealed that collagen did not self-assemble into fibers when mixed with either additive in drop-cast (**Figure S2a-b**) or 3D-printed (**Figure S2c-d**) gels. Instead, large fragments of reflective material, which were likely aggregates of collagen and the additives, were observed. Similar aggregation has been reported for mixtures of collagen with polyethylene glycol or hyaluronic acid ^20^. These results suggest that collagen self-assembly is adversely affected by the charged Laponite particles and the Pluronic micelles. These possibilities are further supported by measurements of the shear storage moduli for the inks: collagen-Laponite and collagen-Pluronic are significantly different than pure collagen, Laponite, or Pluronic (**Figure S3**).

We therefore searched for an alternative material to improve shape retention and printability without disrupting collagen self-assembly. Matrigel is a gelatinous mixture of basement membrane proteins ^41, 42^, primarily laminin (∼60%), type IV collagen (∼30%), and entactin (∼8%). Matrigel is compatible with collagen fibrillogenesis ^2^ and also gels at similar temperatures to collagen (∼37°C), as shown by measurements of its temperature-dependent G′ and G′′ values (**Figure S3**). We therefore mixed Matrigel with collagen and examined the shape retention and printability of the resulting inks. CRM images of drop-cast (**Figure 1d**) and 3D-printed (**Figure 1e**) collagen-Matrigel samples revealed intact collagen fiber networks. In addition, collagen-Matrigel inks were found to have good printability and shape retention (**Figure S1**) and could be 3D-printed into narrow (∼600 µm) lines (**Figure 1f and g**). We therefore further explored the use of collagen-Matrigel inks for 3D-printing.

We evaluated collagen fiber anisotropy in drop-cast and 3D-printed inks by quantifying the fraction of aligned collagen fibers. We found that the alignment fraction of drop-cast (27.9 ± 0.79%) and 3D-printed (27.9 ± 1.4%) pure collagen inks was identical, indicating that 3D-printing did not alter the anisotropy of collagen fibers (**Figure 1h**). Increasing collagen concentration (1.6 mg/ml; 27.9 ± 1.1% or 2.4 mg/ml; 28.2 ± 1.1%) did not alter collagen fiber anisotropy (**Figure S4**). Similar results were observed with collagen-Laponite (**Figure S2e**) and collagen-Pluronic (**Figure S2f**) inks. In contrast, fiber alignment was significantly higher in 3D-printed rectangles (31.5 ± 1.1%) and lines (38.1 ± 0.87%) than in drop-cast (27.0 ± 0.18%) collagen-Matrigel samples (**Figure 1i**). To evaluate the spatial distribution of collagen anisotropy in 3D-printed samples, we acquired CRM images at varying x and y positions and generated alignment fraction heat maps (**Figure S5**). These heat maps qualitatively show that 3D-printed collagen-Matrigel samples had regions of higher alignment than 3D-printed collagen samples and were more uniformly aligned than 3D-printed collagen-Laponite or collagen-Pluronic samples. Alignment fraction did not vary as a function of sample depth (**Figure S5**).

To identify the mechanism by which collagen is aligned during 3D microextrusion printing, we investigated why anisotropy is generated in collagen-Matrigel inks but not pure collagen. One possible explanation is that molecular crowding in collagen-Matrigel improves anisotropy in 3D-printed samples by increasing the molecular weight of collagen assemblies prior to extrusion. To test this hypothesis, we incorporated different concentrations of the molecular crowders Ficoll 70 and Ficoll 400 into collagen inks that were then drop-cast or 3D-printed. CRM images of 3D-printed collagen-Ficoll 400 hydrogels showed isotropic networks at Ficoll concentrations of 6 mg/ml (27.3 ± 0.27%) and 12 mg/ml (27.7 ± 1.4%), anisotropic networks at 18 mg/ml (37.5 ± 1.7%), and entangled networks at 24 mg/ml (28.8 ± 2.3%) (**Figure 2a-e**). Ficoll 70 gave similar results (**Figure S6a-e**). Adding Ficoll did not increase collagen anisotropy in drop-cast samples (**Figure S6f-n**). These results suggest that there is an optimal range for the molecular weight of collagen assemblies required to fabricate anisotropic networks of collagen. Below this range, collagen assemblies are either too small to be aligned or not stable enough to be extruded without mechanical disruption. Above this range, collagen assemblies are either too large to print without clogging the nozzle or are extruded as isotropic entangled collagen networks.

**Figure 2.**
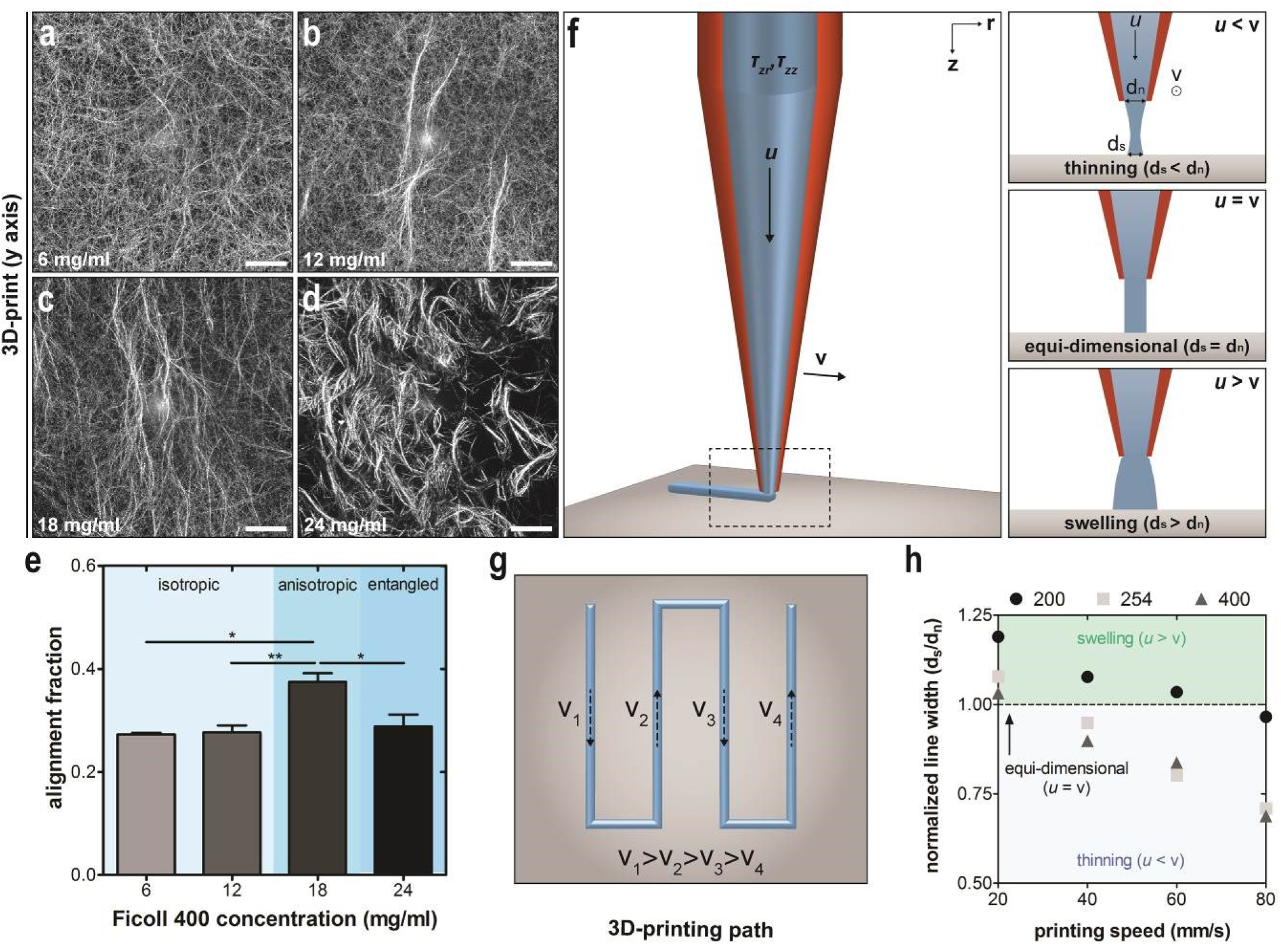
Identifying the mechanism for collagen fiber anisotropy in 3D-printed collagen inks. CRM images of 3D-printed collagen inks containing Ficoll 400 concentrations of a) 6 mg/ml, b) 12 mg/ml, c) 18 mg/ml, or d) 24 mg/ml. All CRM images represent the maximum-intensity z-projection of a 30-µm z-stack. e) Alignment fraction of 3D-printed collagen-Ficoll 400 inks. Collagen-Ficoll samples were printed using a 254-µm-diameter conical nozzle at a printing speed of 40 mm/s. Scale bars = 50 µm. f) Schematic of conical nozzle during 3D-printing and g) 3D-printing path used to identify different printing regimes shown in panel f. h) Normalized width of 3D-printed line as a function of printing speed and nozzle diameter.

During 3D microextrusion printing, hydrogels are subjected to shear and extensional flows in the conical printing nozzle, which may be sufficient to align collagen assemblies in the printing direction. Classical lubrication theory applied to the 254-µm-diameter conical nozzle yields shear and strain rate estimates on the order of 100 s^-1^. This order of magnitude is consistent with shear rates previously reported to align type I collagen ^16, 43^. Additionally, we investigated three printing regimes that are based on the relative speeds of ink extrusion (*u*) and nozzle translation (*v*), which include thinning (*u* < *v*), equi-dimensional (*u* = *v*), and swelling (*u* > *v*) (**Figure 2f**).^44^ We demonstrated that all three regimes could be achieved by 3D-printing collagen-Matrigel inks at speeds ranging from 20 to 80 mm/s with nozzle diameters ranging from 200 to 400 µm (**Figure 2g and h**). These data show that the collagen-Matrigel ink can experience stretching or accumulation after exiting the nozzle, which may impact anisotropy in the printed sample.

Given the suspected role of shear and extensional flows in aligning collagen assemblies during printing, we next proceeded to investigate the range over which collagen fiber anisotropy and geometry can be tuned. We 3D-printed collagen-Matrigel inks in which we varied Matrigel protein concentration (**Figure 3a-c**). The average collagen fiber diameter was unaffected by changing Matrigel concentration (**Figure S7a**). However, the average length of collagen fiber bundles was significantly higher at a Matrigel concentration of 3 mg/ml (117 ± 11 µm) than at higher concentrations (4.5 mg/ml, 68.4 ± 5.8 µm; 6.0 mg/ml, 47.5 ± 2.4 µm) (**Figure S7b**). These results are consistent with findings that the concentration of Matrigel can impact collagen fiber morphology ^45^. We found that collagen alignment remained constant as Matrigel protein concentration varied from 3 mg/ml (28.36 ± 0.76%) to 6 mg/ml (26.68 ± 0.46%) (**Figure 3d**).

**Figure 3.**
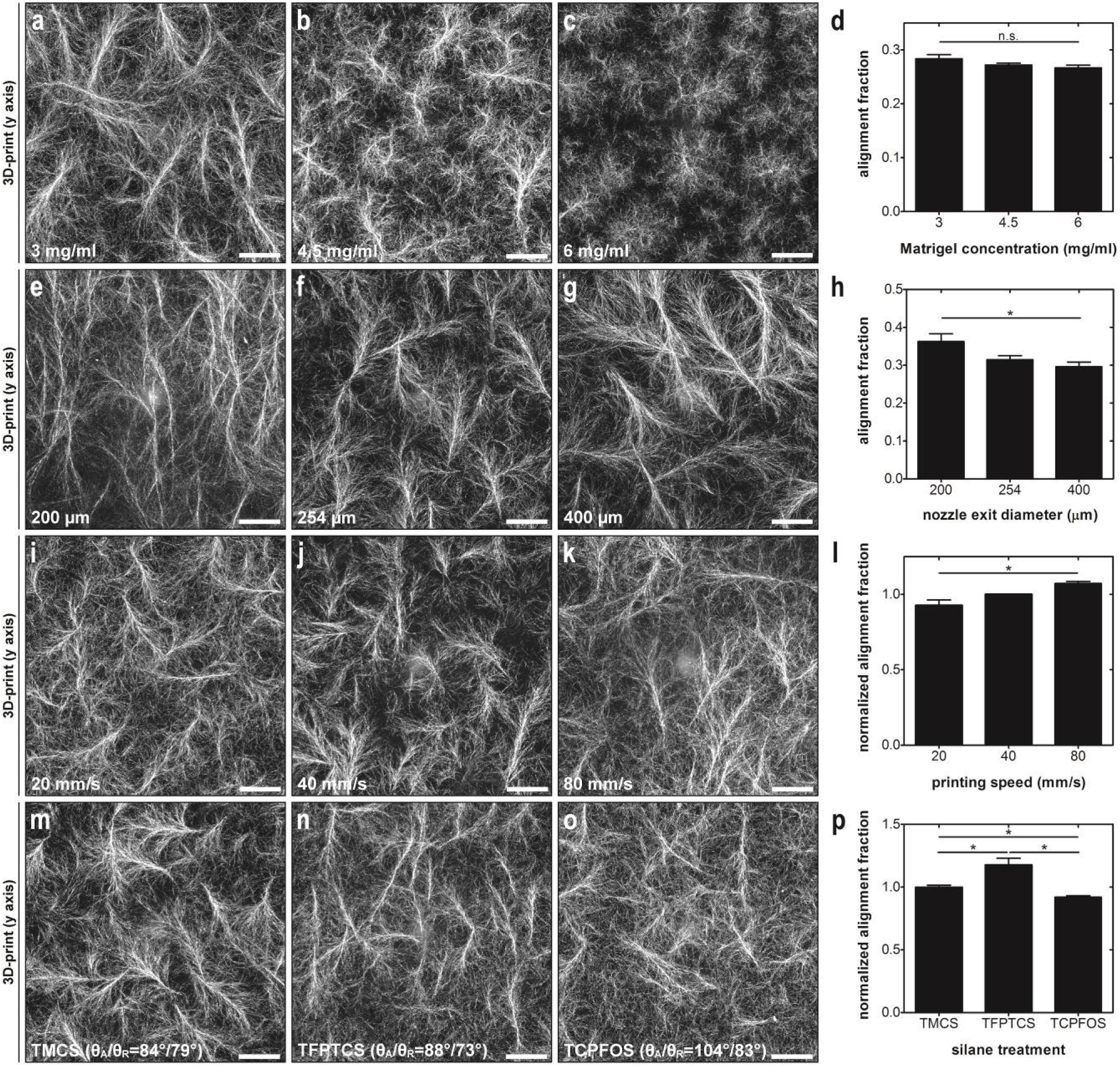
Tuning collagen fiber geometry and anisotropy. CRM images of 3D-printed collagen-Matrigel samples with Matrigel concentrations of a) 3.0 mg/ml, b) 4.5 mg/ml, or c) 6.0 mg/ml. d) Alignment fraction as a function of Matrigel concentration. CRM images of collagen-Matrigel inks printed using conical nozzles with a diameter of e) 200 µm, f) 254 µm, or g) 400 µm. h) Alignment fraction of collagen-Matrigel inks printed using different nozzle diameters. CRM images of collagen-Matrigel inks printed using a 254-µm-diameter nozzle at printing speeds of i) 20 mm/s, j) 40 mm/s, or k) 80 mm/s. l) Normalized alignment fraction of collagen-Matrigel inks 3D-printed at different speeds. CRM images of collagen-Matrigel inks printed using a 254-µm-diameter nozzle onto glass coverslips functionalized with m) TMCS, n) TFPTCS, and o) TCPFOS. Advancing and receding water contact angles are provided at the bottom of each image. p) Normalized alignment fraction of collagen-Matrigel inks 3D-printed onto different surface chemistries. Scale bars on CRM images = 50 µm. All CRM images represent the maximum-intensity z-projection of a 30-µm z-stack. A Matrigel concentration of ∼5.0 mg/ml was used for all collagen-Matrigel samples in panels e-p.

To investigate the effects of nozzle diameter and printing speed on collagen fiber anisotropy, we 3D-printed collagen-Matrigel inks using conical nozzles with different exit diameters at different speeds. Collagen anisotropy increased as nozzle diameter decreased (**Figure 3e-h**), which is consistent with the suspected role of shear and extensional flows in aligning collagen during printing. Similarly, we found that the alignment fraction increased with increasing printing speed (**Figure 3i-l**).

We observed that collagen-Matrigel inks had reduced surface wetting as compared to pure collagen inks (**Figure S8a**), which suggested that surface chemistry might play a role in dictating the anisotropy of collagen fibers. To test this hypothesis, we used different silane treatments to modify the hydrophobicity of the glass substratum on which the collagen inks were printed, which was evaluated using advancing (θ_A_) and receding (θ_R_) water contact angles. Collagen-Matrigel inks were 3D-printed onto hydrophobic trimethylchlorosilane (TMCS) (θ_A_/θ_R_ = 84°/79°), 3,3,3-trifluoropropyl-trichlorosilane (TFPTCS) (θ_A_/θ_R_ = 88°/73°), and trichloro(1H,1H,2H,2H-perfluorooctyl)silane (TCPFOS) (θ_A_/θ_R_ = 104°/83°) -treated glass, while methoxy-poly(ethylene glycol)-silane (PEG-silane) (θ_A_/θ_R_ = 35°/21°) was used as a hydrophilic control (**Figure S8b**). Advancing contact angle measurements showed that collagen-Matrigel inks became increasingly dewetting as substratum hydrophobicity increased (**Figure S8c**). We quantified a normalized alignment fraction, which represents the alignment fraction on silane-treated glass relative to a control from the same ink printed onto untreated glass. We found that collagen anisotropy is maximized at intermediate substratum hydrophobicity (TFPTCS) (**Figure 3m-p**). We conclude that there is an ideal range for substratum hydrophobicity that maximizes alignment. Below this range the ink spreads after exiting the nozzle (**Figure 3m**), and above this range the ink coalesces, which also reduces alignment (**Figure 3o**). Taken together, these data reveal that anisotropy can be tuned by modulating shear and extensional flows during printing as well as the hydrophobicity of the underlying substratum.

Controlling these parameters allowed us to spatially pattern collagen fiber alignment and geometry. By 3D-printing collagen-Ficoll and collagen-Matrigel inks into a fan-shaped pattern (**Figure 4a**), we achieved multidirectional collagen fiber alignment (**Figure 4b-d**). A lower concentration of Matrigel (3 mg/ml) was needed to obtain alignment in the fan-shaped pattern, which suggests that the optimal Matrigel concentration depends on the printed geometry. To demonstrate control of shape retention and printability, we 3D-printed an outline of a shield using collagen-Matrigel (**Figure 4e**). We found that we could spatially control collagen fiber geometry by printing lines of collagen-Matrigel with a high Matrigel concentration (7.2 mg/ml) and drop-casting low Matrigel concentrations (3 mg/ml) in between (**Figure 4f**). CRM images revealed distinct regions with different collagen fiber geometries that were separated by sharp interfaces (**Figure 4f**). To demonstrate the flexibility of our approach, we simultaneously controlled collagen fiber alignment and geometry by drop-casting pure collagen in between 3D-printed lines of aligned collagen-Matrigel (**Figure 4g**). We observed that collagen fibers were aligned at the interface between these two collagen morphologies, which is consistent with previous studies of collagen-collagen ^46^ and collagen-Matrigel ^18^ interfaces. Moreover, we were able to generate complex alignment patterns by 3D-printing interfaces with both concave (**Figure 4h**) and convex curvature (**Figure 4i**). These results demonstrate that 3D-printing can be used to generate a variety of complex patterns of collagen fiber anisotropy and geometry that cannot be achieved with conventional alignment techniques.

**Figure 4.**
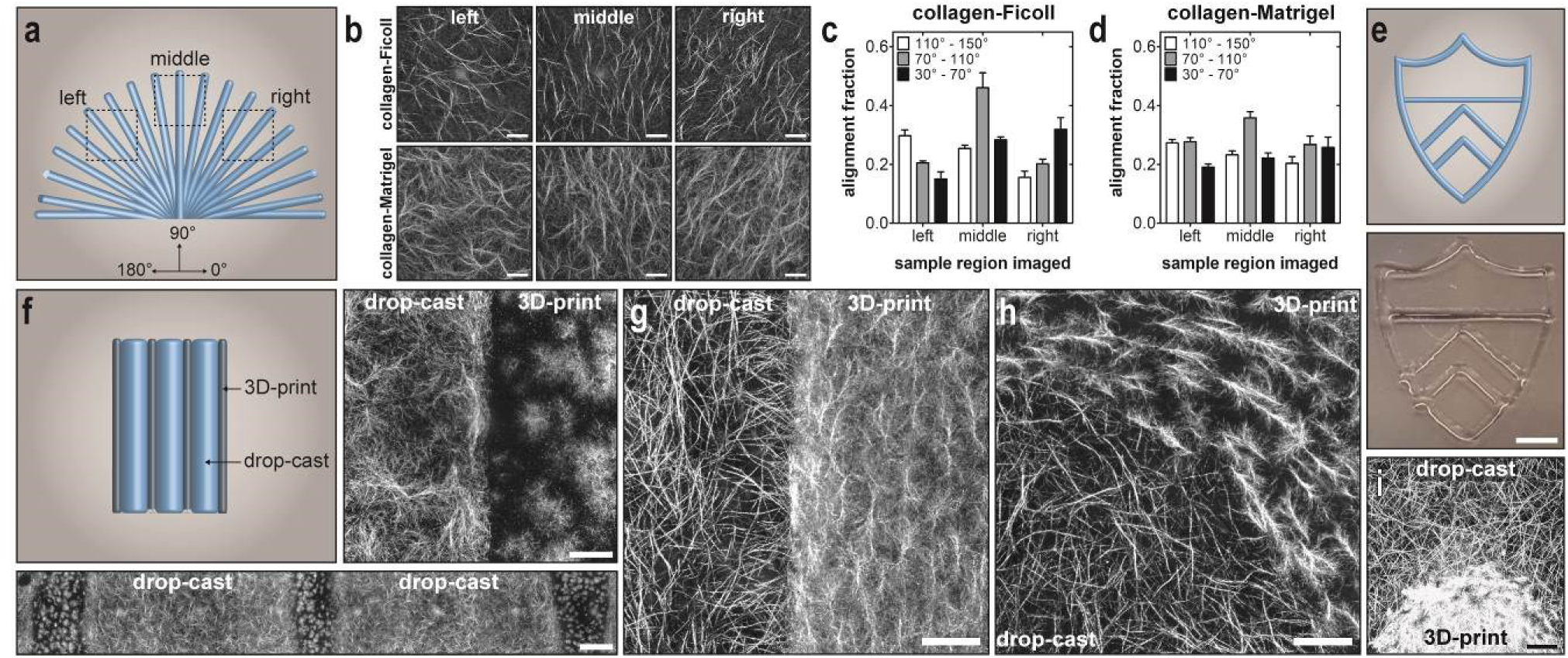
3D-printing patterns of collagen fiber anisotropy and geometry. a) Schematic of 3D-printing path used to demonstrate multidirectional collagen fiber alignment. Boxes indicate the three regions that were imaged. b) CRM images of collagen-Ficoll and collagen-Matrigel inks 3D-printed using the printing path shown in panel a. Scale bars = 100 µm. Alignment fraction of 3D-printed c) collagen-Ficoll and d) collagen-Matrigel samples printed using the printing path in panel a. Ficoll 400 was used at a concentration of 18 mg/ml and Matrigel was used at a concentration of 3 mg/ml. Samples were 3D-printed using a 254-µm-diameter nozzle at a speed of 80 mm/s. e) 3D-printing path of a shield (top) and optical image of 3D-printed collagen-Matrigel ink (bottom; scale bar = 2.5 mm). Matrigel concentration was 6.8 mg/ml and the glass substratum was functionalized with TFPTCS. f) Schematic of 3D-printed lines and drop-cast regions along with CRM images of the interface between drop-cast and 3D-printed collagen-Matrigel regions (right; scale bar = 50 µm) and a slice across the sample consisting of 8 images stitched together (bottom; scale bar = 250 µm). Concentration of Matrigel was 3.0 mg/ml and 7.2 mg/ml for the drop-cast and 3D-printed inks, respectively. CRM images of g) straight, h) concave, and i) convex interfaces between drop-cast collagen and 3D-printed collagen-Matrigel (5.8 mg/ml); scale bars = 100 µm. All CRM images represent the maximum-intensity z-projection of a 30-µm z-stack.

To demonstrate the utility of 3D-printed collagen networks, we cultured MDA-MB-231 human breast cancer cells on top of 3D-printed collagen-Matrigel scaffolds. Based on previous reports that breast cancer cells orient in the direction of collagen fiber alignment ^18^, we hypothesized that cells would orient in the printing direction. Cells were suspended in culture medium and subsequently drop-cast on top of polymerized collagen-Matrigel. After 60 h of incubation, cells were oriented along the direction of collagen fiber alignment for both 3D-printed lines and curved geometries (**Figure 5a and b**). We found that the cell alignment fraction, which represents the fraction of cells oriented within 20° of the fiber alignment direction, was significantly higher for cells cultured on anisotropic (44.81 ± 1.93%) 3D-printed collagen-Matrigel networks as compared to isotropic (27.27 ± 3.16%) networks (**Figure 5c**). In addition, we incorporated human breast cancer cells into unpolymerized collagen, which was subsequently drop-cast on top of and in between 3D-printed lines of collagen-Matrigel (as in **Figure 4g**). After 24 h of incubation, the cells were oriented along collagen fibers that were aligned perpendicular to the 3D-printed collagen-Matrigel interface (**Figure 5d**) and appeared to migrate towards the interface (**Figure 5e**), which is consistent with previously reported results ^18^. We found that the cell alignment fraction within 100 µm of the interface (33.25 ± 5.52%), where collagen fibers are aligned, was higher than in isotropic collagen networks (19.60 ± 1.79%). We also incorporated mammary epithelial cell clusters into the collagen-Matrigel ink and microextrusion-printed this bioink into straight lines on a glass substratum. After printing, the cell-laden collagen-Matrigel construct was cured at 37°C, submerged in culture medium supplemented with 5 ng/ml hepatocyte growth factor (HGF), and cultured for two days. Phase-contrast images of the cell clusters were acquired immediately after curing (**Figure 5g**) as well as after one (**Figure 5h**) and two (**Figure 5i**) days in culture. These data reveal that mammary epithelial cell clusters can be cultured in 3D-printed collagen-Matrigel networks, and that the clusters extend protrusions in the direction of collagen fiber alignment. As the number of days in culture increased, the circularity of clusters decreased and the orientation of the clusters approached the orientation of the printing direction (0°) (**Figure 5j**). These results demonstrate that 3D-printed collagen networks can be used to study a wide variety of interactions between cells and aligned networks of collagen fibers.

**Figure 5.**
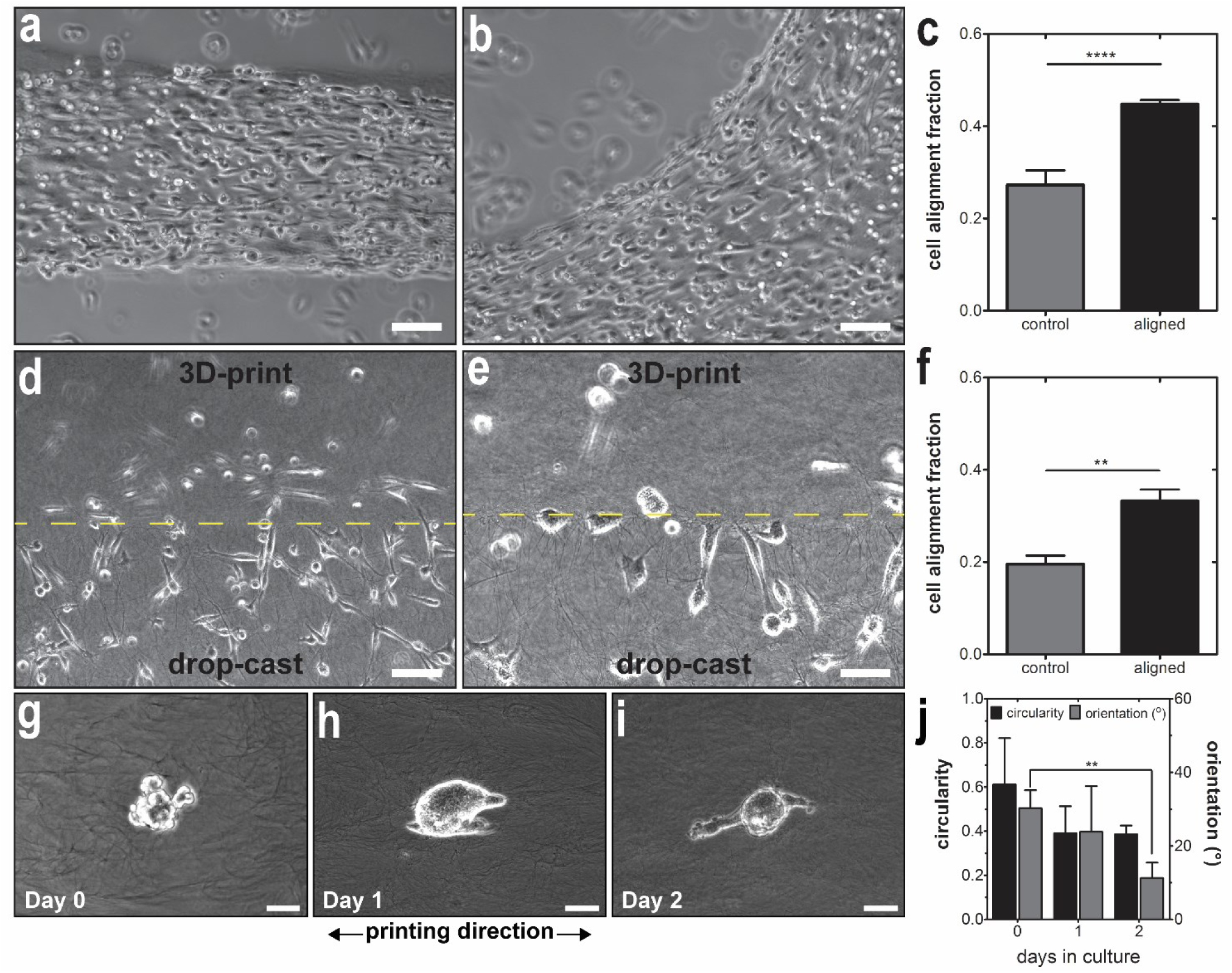
Culturing cells on top of and within 3D-printed networks of collagen. Phase-contrast images of MDA-MB-231 human breast cancer cells cultured on top of collagen-Matrigel inks 3D-printed in a a) straight line and b) curved geometry. c) Alignment fraction of cells on top of 3D-printed lines. The control consists of cells cultured on top of collagen-Matrigel networks with isotropic collagen fiber orientation. Cells were cultured for 60 h before image acquisition. d) Low magnification and e) high magnification phase-contrast image of the interface, denoted by dashed line, between a line of 3D-printed collagen-Matrigel and drop-cast collagen. Human breast cancer cells were suspended in collagen, which was drop-cast on top of and in between 3D-printed lines of collagen-Matrigel. Cells were subsequently cultured for 24 h before image acquisition. f) Alignment fraction of cells in the drop-cast collagen phase within ∼100 µm of the interface. Binary images were used to determine cell orientation near the interface. The control consists of cells cultured within collagen with isotropic fiber orientation. Phase-contrast images of mammary epithelial cell clusters g) 0, h) 1, and i) 2 days after 3D-microextrusion printing in a collagen-Matrigel ink with a volumetric ratio of 7 parts collagen and 3 parts Matrigel. j) Circularity and Feret diameter orientation of 3D-printed epithelial cell clusters. Data in plots c and f represent the average of 5 replicates while data in plot j represents the average of 3 tissues. Error bars on all plots represent standard deviation. A two-tailed *p*-value was used for comparison with the control in plots c and f. Scale bars = 100 µm for a, b, and d and 50 µm for e, g, h, and i.

While collagen-Matrigel inks enable the fabrication of anisotropic networks of collagen fibers, the batch-to-batch variability and high cost of Matrigel preclude its use in clinical settings. This motivates the need for a similar gelatinous material that is compatible with collagen fibrillogenesis and that has desirable viscoelastic properties for 3D microextrusion printing. A synthetic formulation of Matrigel might be useful for improving printability and shape retention, which would allow for the fabrication of 3D cell-laden constructs, while preserving native collagen fiber structure.

## Conclusions

Here, we showed that 3D microextrusion printing of collagen-Matrigel inks can be used to fabricate cell-laden anisotropic networks of type I collagen fibers. By tuning collagen self-assembly conditions and printing parameters, we demonstrated that collagen fiber geometry and anisotropy can be spatially patterned. Our results suggest that shear and extensional forces generated during 3D printing are responsible for aligning collagen and that the size of collagen assemblies during printing, which is dictated by molecular crowding in the ink, can be used to modulate anisotropy in 3D-printed networks of collagen. Moreover, we showed that cells cultured on top of 3D-printed collagen networks orient in the direction of collagen fiber alignment. We also demonstrated that collagen-Matrigel inks could be used to bioprint cell-laden constructs, wherein aligned networks of collagen surround fully embedded epithelial cell clusters. Microextrusion printed cell-laden anisotropic networks of collagen fibers have broad utility in fields ranging from developmental biology to tissue engineering and regenerative medicine.

## Author contributions

B.A.N., P.-T.B., and C.M.N. designed experiments; B.A.N. and P.-T.B. performed modeling calculations; B.A.N. performed experiments, completed data analysis, and prepared figures; B.A.N., P.-T.B., and C.M.N. wrote the manuscript.

## Acknowledgements

We thank J. Tien and members of the Tissue Morphodynamics Group for insightful discussions. We also thank G. Laevsky and the Molecular Biology Confocal Microscopy Facility (Princeton University) for assistance with CRM imaging, and L. Loo for access to the contact angle goniometer and cleanroom. This work was supported in part by grants from the NIH (HL118532, HL120142, CA187692), the NSF (CMMI-1435853), the David & Lucile Packard Foundation, the Camille & Henry Dreyfus Foundation, and Princeton University’s Project X Fund. B.A.N. was supported in part by a postgraduate scholarship-doctoral (PGS-D) from the Natural Sciences and Engineering Research Council of Canada. C.M.N. was supported in part by a Faculty Scholars Award from the Howard Hughes Medical Institute.

## Abbreviations

2D: two-dimensional
3D: three-dimensional
CRM: confocal reflection microscopy
ECM: extracellular matrix

