## Supplemental Material for "Microextrusion Printing Cell-Laden Networks of Type I Collagen with Patterned Anisotropy and Geometry"

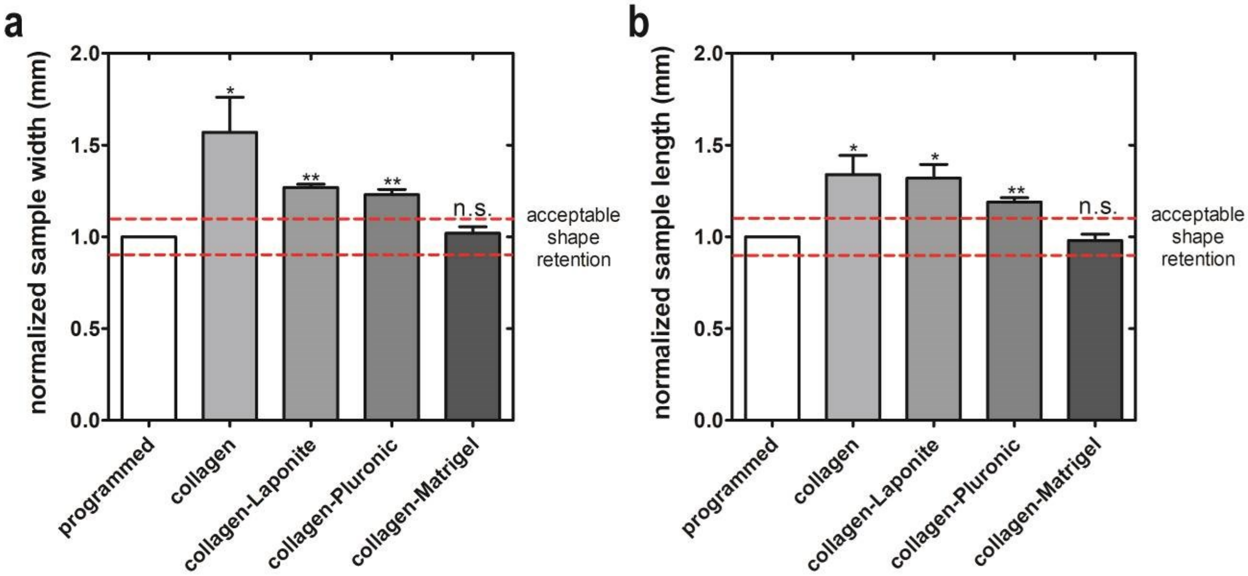


**Figure S1. Shape retention of collagen inks.** Normalized sample a) width and b) length of rectangular geometries 3D-printed with collagen ink formulations. "Programmed" represents the dimensions of the printing path, while normalized dimensions are calculated by taking the ratio of measured dimensions relative to the programmed dimensions. Dashed red lines represent the criteria for acceptable shape retention (±10% of programmed dimensions).

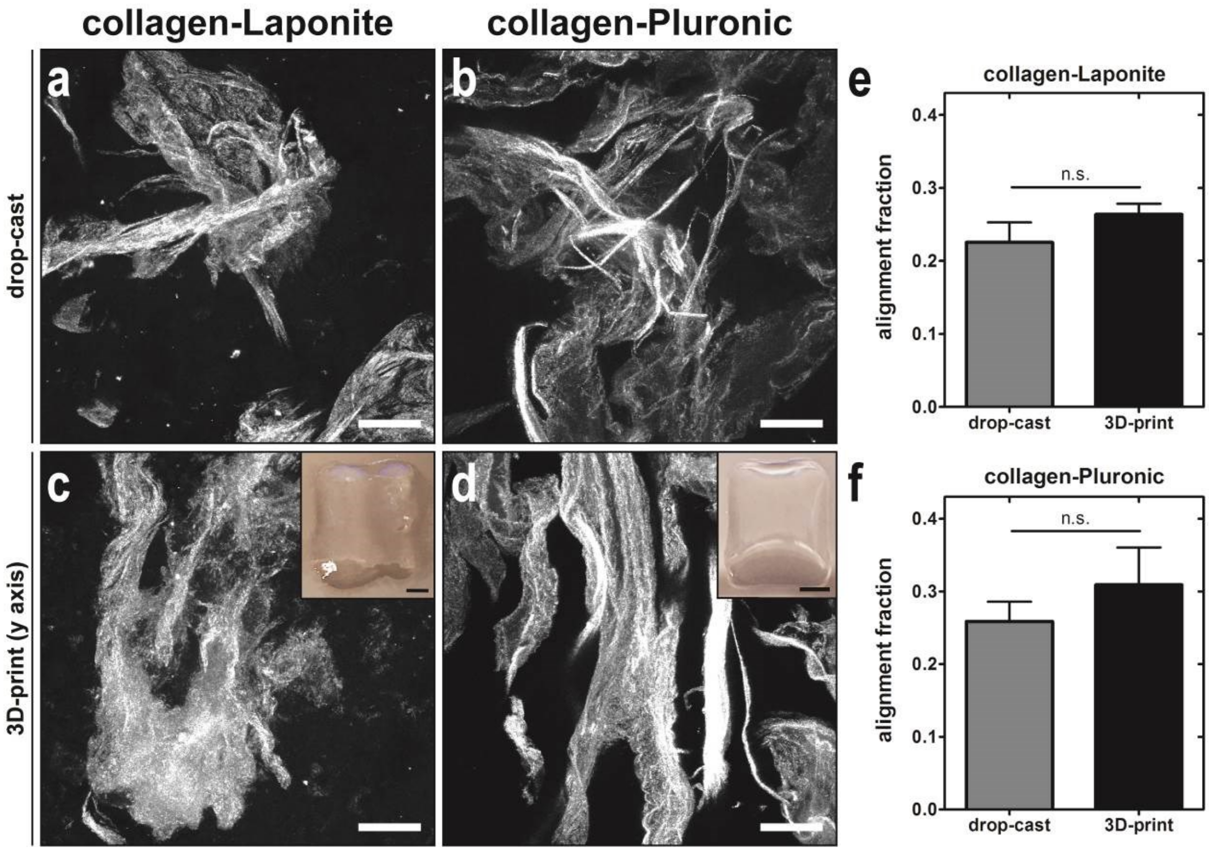


**Figure S2. Morphology and anisotropy of collagen-Laponite and collagen-Pluronic inks.** CRM images of drop-cast a) collagen-Laponite and b) collagen-Pluronic and 3D-printed c) collagen-Laponite and d) collagen-Pluronic. All CRM images represent the maximum-intensity z-projection of a single 30-µm-thick z-stack. Optical images are representative 3D-printed rectangular samples. Samples were 3D-printed using a 254-µm-diameter conical nozzle at a printing speed of 40 mm/s. Scale bars on CRM images represent 50 µm, and scale bars on optical images represent 2.5 mm. Average alignment fraction of drop-cast and 3D-printed samples of e) collagen-Laponite and f) collagen-Pluronic.

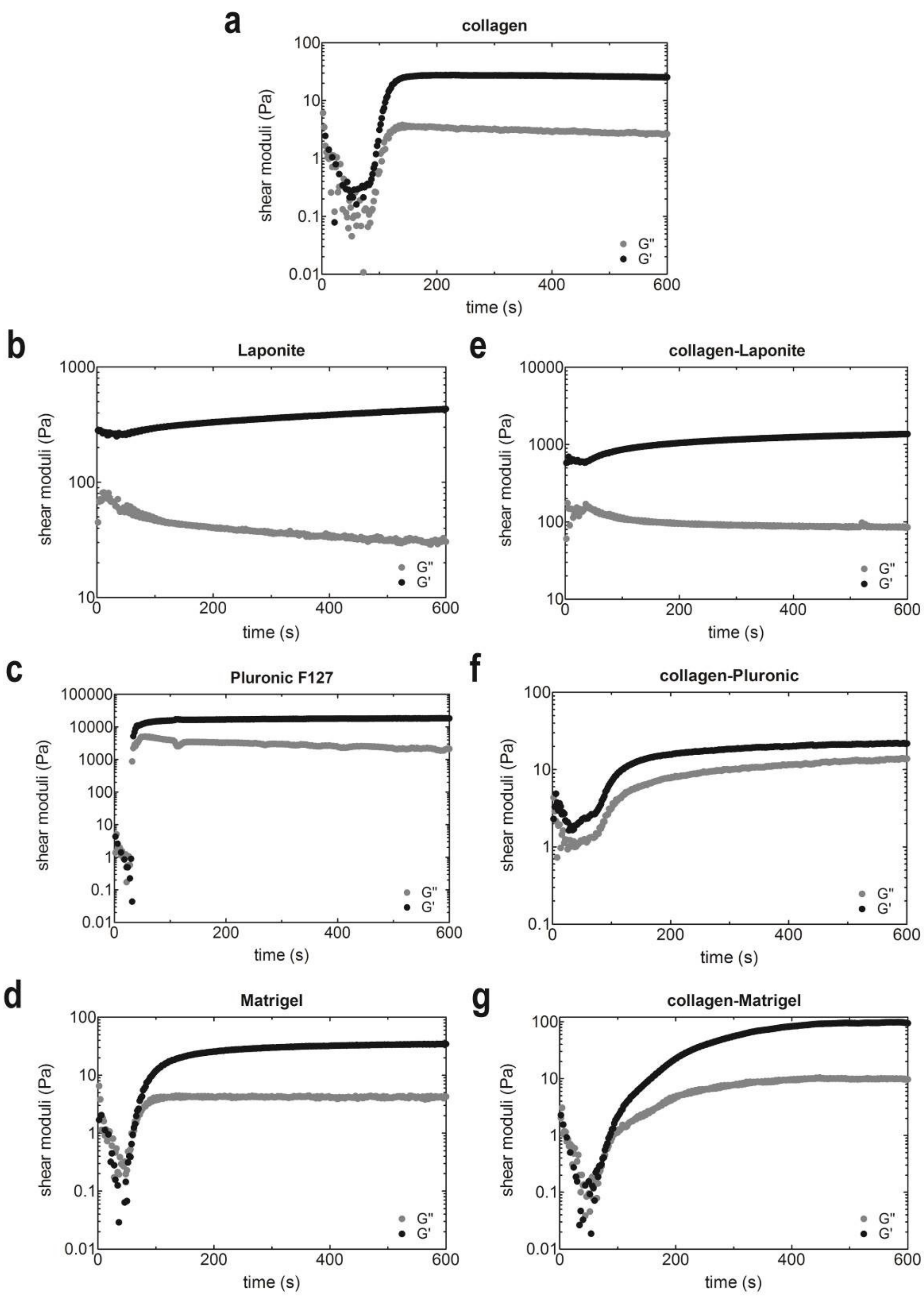


**Figure S3. Shear storage and loss moduli of collagen inks.** Representative temperature-dependent storage and loss moduli for a) 0.8 mg/ml type I collagen, b) 3 mg/ml Laponite, c) 250

mg/ml Pluronic F127, d) 8.2 mg/ml Matrigel, e) collagen-Laponite, f) collagen-Pluronic, or g) collagen-Matrigel. Data from first replicate is plotted; outliers at early time points were excluded.

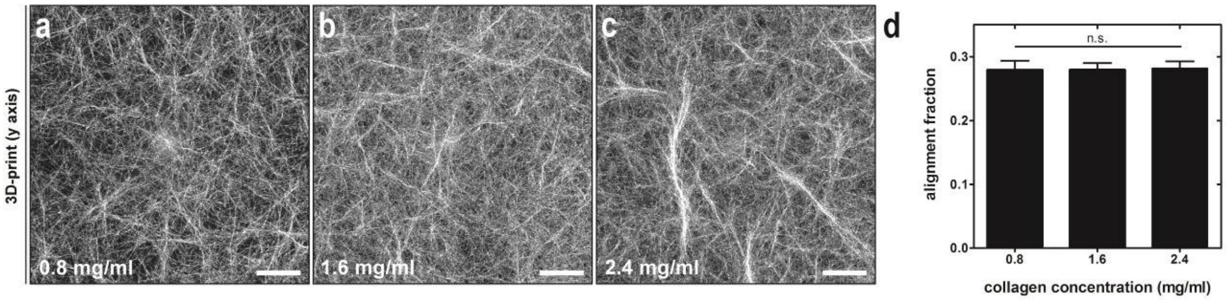


**Figure S4. 3D-printing different concentrations of type I collagen.** CRM images of type I collagen inks with concentrations of a) 0.8 mg/ml, b) 1.6 mg/ml, or c) 2.4 mg/ml. d) Alignment fraction of 3D-printed collagen samples with different collagen concentrations. Scale bars = 50 µm. All CRM images represent the maximum-intensity z-projection of a 30-µm-thick z-stack.

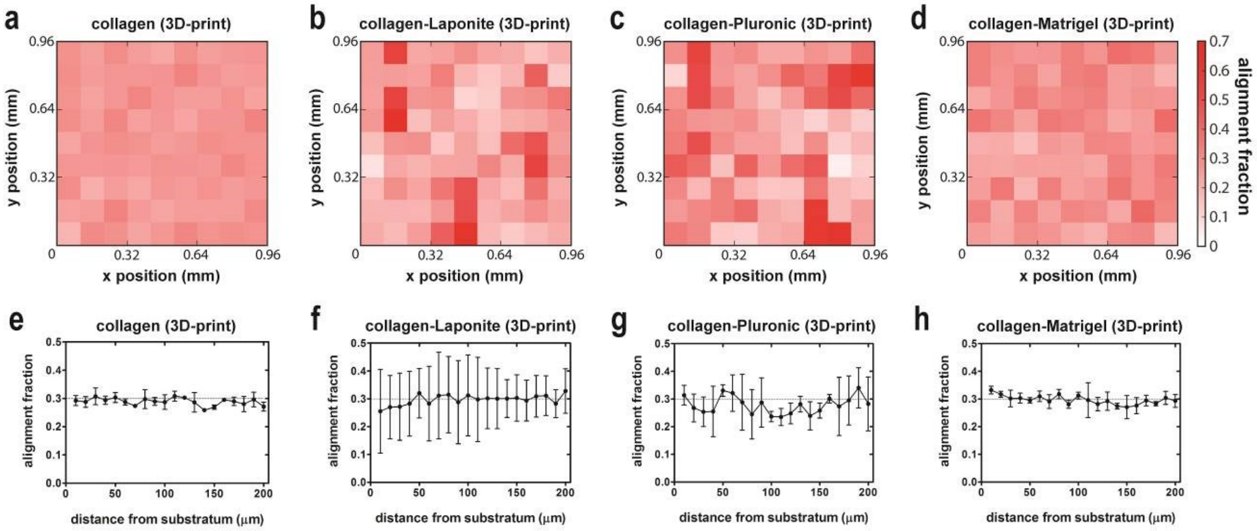


**Figure S5. Quantifying collagen fiber alignment in 3D-printed samples.** Alignment fraction heat map for a representative sample of 3D-printed a) type I collagen, b) collagen-Laponite, c) collagen-Pluronic, or d) collagen-Matrigel. Alignment fraction as a function of depth for a representative sample of 3D-printed e) type I collagen, f) collagen-Laponite, g) collagen-Pluronic, or h) collagen-Matrigel. A dotted reference line is included at an alignment fraction of

0.3. All drop-cast and 3D-printed samples were incubated at ~0°C for 1 h after neutralization, and the collagen concentration was 0.8 mg/ml for all samples. All samples were printed using a 254-μm-diameter nozzle at a printing speed of 40 mm/s.

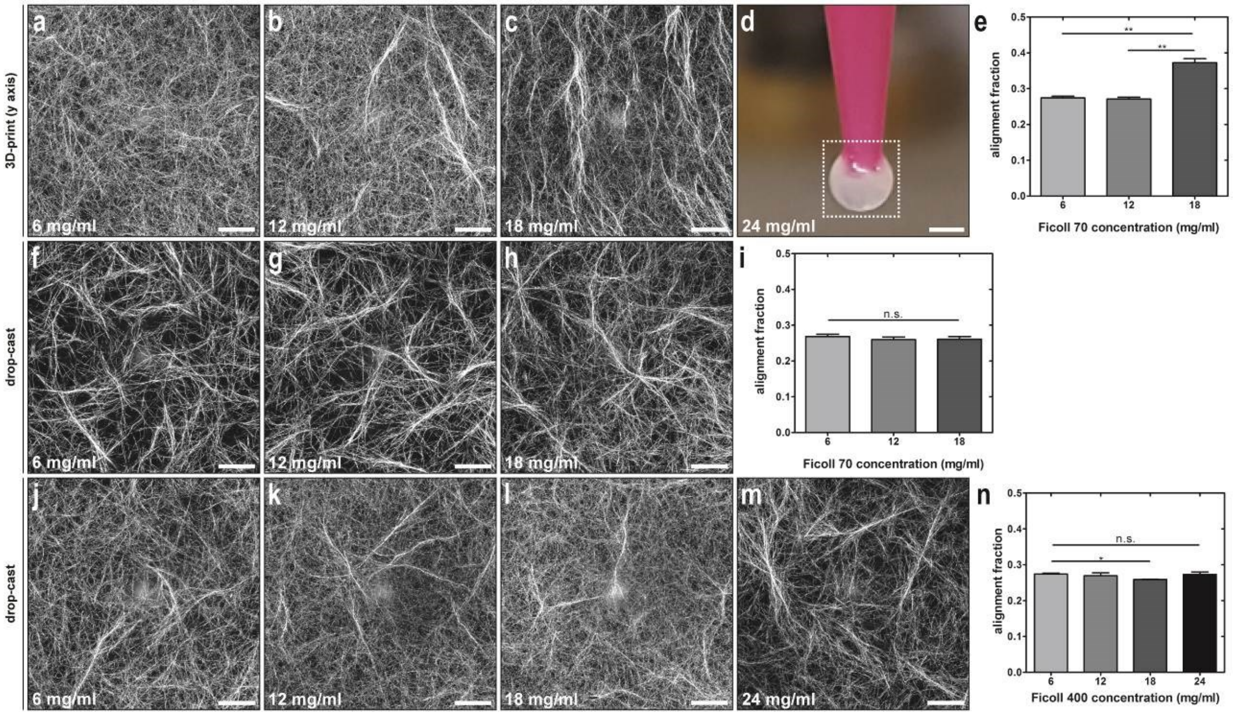


**Figure S6. 3D-printed and drop-cast collagen-Ficoll inks.** CRM images of 3D-printed collagen inks containing Ficoll 70 concentrations of a) 6 mg/ml, b) 12 mg/ml, or c) 18 mg/ml. d) Optical image of the 3D-printing nozzle used to print a collagen-Ficoll 70 ink with a Ficoll 70 concentration of 24 mg/ml. The image shows polymerized collagen blocking the nozzle in the region enclosed in white dashed lines. e) Alignment fraction for 3D-printed collagen-Ficoll 70 inks. CRM images of drop-cast collagen inks containing Ficoll 70 concentrations of f) 6 mg/ml, g) 12 mg/ml, or h) 18 mg/ml. i) Alignment fraction for drop-cast collagen-Ficoll 70 inks. CRM images of drop-cast collagen inks containing Ficoll 400 concentrations of j) 6 mg/ml, k) 12 mg/ml, l) 18 mg/ml, or m) 24 mg/ml. n) Alignment fraction for drop-cast collagen-Ficoll 400 inks. Scale bars = 50 μm. All CRM images represent the maximum-intensity z-projection of a 30-μm z-stack.

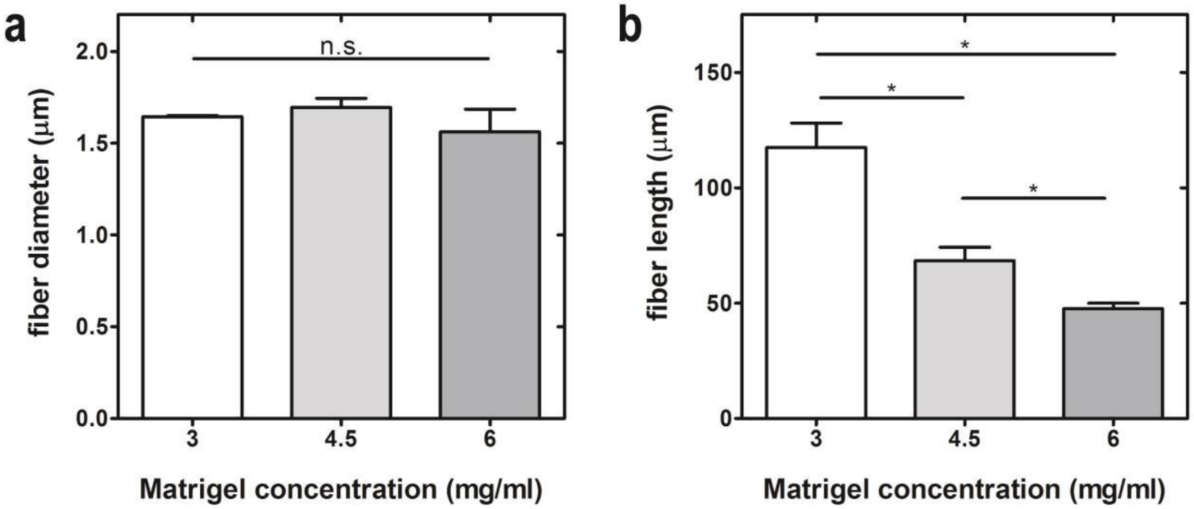


**Figure S7. Collagen fiber diameter and length as a function of Matrigel protein concentration.** Average a) collagen fiber diameter and b) length of collagen fiber bundles in 3D-printed collagen-Matrigel samples with Matrigel protein concentrations of 3 mg/ml, 4.5 mg/ml, or 6 mg/ml. All samples were 3D-printed using a 254-µm-diameter nozzle at a printing speed of 40 mm/s.

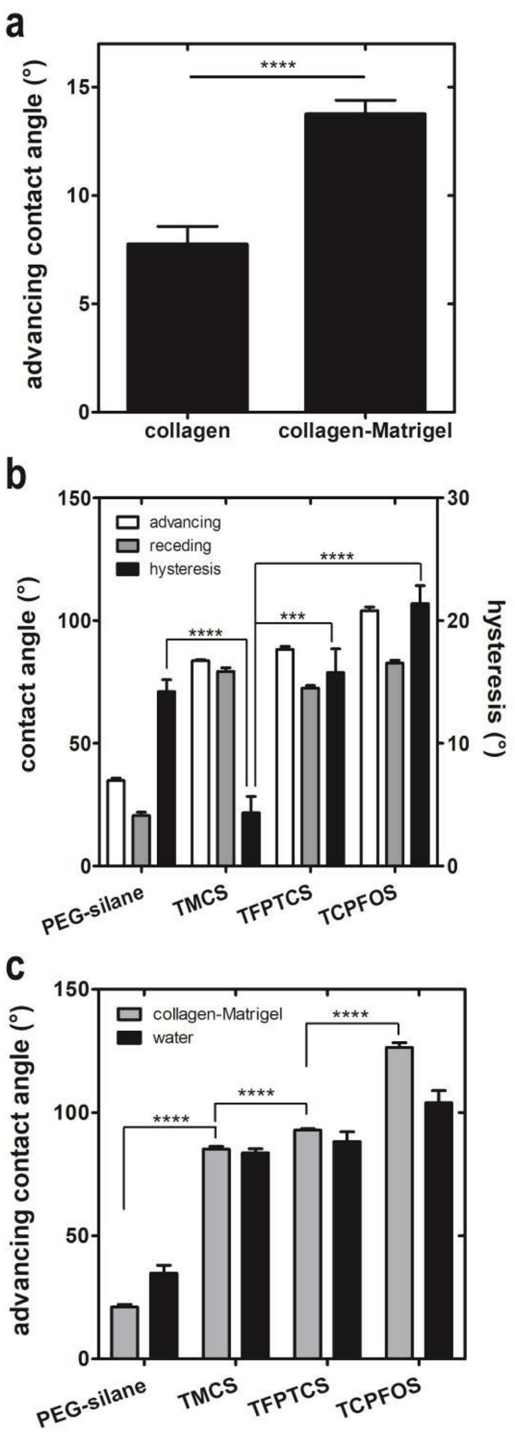


**Figure S8. Contact angle measurements of collagen inks.** a) Advancing contact angle measurements for collagen and collagen-Matrigel on untreated glass. b) Advancing and receding contact angles for water on silanized glass. c) Advancing contact angle measurements for collagen-Matrigel and water on silanized glass. The average of 10 experimental measurements is plotted for all contact angle data, and error bars represent standard error of the mean. Two-sided *p*-values were use
